# Broad substrate-specific phosphorylation events are associated with the initial stage of plant cell wall recognition in *Neurospora crassa*

**DOI:** 10.1101/711085

**Authors:** Maria Augusta Crivelente Horta, Nils Thieme, Yuqian Gao, Kristin E. Burnum-Johnson, Carrie D. Nicora, Marina A. Gritsenko, Mary S. Lipton, Karthikeyan Mohanraj, Leandro José de Assis, Liangcai Lin, Chaoguang Tian, Gerhard H. Braus, Katherine A. Borkovich, Monika Schmoll, Luis F. Larrondo, Areejit Samal, Gustavo H. Goldman, J. Philipp Benz

## Abstract

Fungal plant cell wall degradation processes are governed by complex regulatory mechanisms, allowing the organisms to adapt their metabolic program with high specificity to the available substrates. While the uptake of representative plant cell wall mono- and disaccharides is known to induce specific transcriptional and translational responses, the processes related to early signal reception and transduction remain largely unkown. A fast and reversible way of signal transmission are post-translational protein modifications, such as phosphorylations, which could initiate rapid adaptations of the fungal metabolism to a new condition. To elucidate how changes in the initial substrate recognition phase of *Neurospora crassa* affect the global phosphorylation pattern, phospho-proteomics was performed after a short (2 minutes) induction period with several plant cell wall-related mono- and disaccharides. The MS/MS-based peptide analysis revealed large-scale substrate-specific protein phosphorylation and de-phosphorylations. Using the proteins identified by MS/MS, a protein-protein-interaction (PPI) network was constructed. The variance in phosphorylation of a large number of kinases, phosphatases and transcription factors indicate the participation of many known signaling pathways, including circadian responses, two-component regulatory systems, MAP kinases as well as the cAMP-dependent and heterotrimeric G-protein pathways. Adenylate cyclase, a key component of the cAMP pathway, was identified as a potential hub for carbon source-specific differential protein interactions. In addition, four phosphorylated F-Box proteins were identified, two of which, Fbx-19 and Fbx-22, were found to be involved in carbon catabolite repression responses. Overall, these results provide unprecedented and detailed insights into a so far less well known stage of the fungal response to environmental cues and allow to better elucidate the molecular mechanisms of sensory perception and signal transduction during plant cell wall degradation.

## 1. Introduction

Fungi are cell factories with important functions for the bioeconomy since they often exhibit superior metabolic and secretory capabilities compared to bacterial and yeast-based production systems (Meyer et al., 2016). Furthermore, fungi are highly versatile in their ability to adjust their metabolism to diverse substrates. In the case of fungal utilization of complex plant biomass, the specific perception of the individual plant cell wall components is a necessary pre-requisite to adapt enzyme production and intracellular metabolism. These recognition reactions trigger a sophisticated series of intracellular signaling cascades in order to control and coordinate the cellular response. The fungal perception of plant cell wall polysaccharides in the presence of plant biomass is initiated by starvation-triggered de-repression of a large number of genes, leading to the production of low amounts of scouting enzymes responsible for plant cell wall depolymerization (van Munster et al., 2014). The resulting monomeric and/or small oligomeric soluble sugars are transferred into the cell via membrane-embedded transporters. Then these sugars (or their metabolic derivatives) can function as signaling molecules that activate signal cascades to promote specific metabolic adaptations (see e.g. Glass et al., 2013 and references therein). The recognition signals are responsible for the transcription and translation of specific genes, such as sugar transporters and enzymes necessary to depolymerize and catabolize the corresponding polysaccharide. However, the signaling reactions that occur between the first substrate contact and the gene activation by transcription factors are poorly described, making it difficult to predict good targets for strain engineering.

Reversible modifications in protein phosphorylation mediated by protein kinases or phosphatases represents a mechanism that is frequently employed by eukaryotic cells to transmit environmental cues and initiate signal transduction pathways, such as for transcription factor activation (Whitmarsh and Davis, 2000), cellular localization, protein stability, protein-protein interactions, DNA binding, and enzymatic activity (Breitkreutz et al., 2010; Pawson, 2007). Most cellular processes are in fact regulated by the reversible phosphorylation of proteins on serine (S), threonine (T), and tyrosine residues (Y) (Ficarro et al., 2002; Wart and Unit, 1993), including metabolism, movement, the circadian rhythms and many other (Cohen, 2000; Hurley et al., 2016; Wart and Unit, 1993).

At least 107 serine/threonine protein kinases and more than 30 nucleases and phosphatases are predicted in *Neurospora crassa* (Borkovich et al., 2004; Galagan et al., 2003; Miranda-Saavedra and Barton, 2007; Park et al., 2011), a filamentous ascomycete that has been used as a model system for genetics for many decades and has available a fully sequenced and well-annotated genome (Borkovich et al., 2004). Since *N. crassa* is furthermore a saprotroph growing robustly on complex lignocellulosic material (Seibert et al., 2016; Tian et al., 2009), we used this fungus in the current study to analyze to what extent post-translational protein modifications by phosphorylation are involved in metabolism-related signaling cascades leading to the perception of the individual polysaccharides (cellulose, hemicelluloses and pectin) within complex plant biomass. Little information is available to this end, however regarding the very early time points of the perception event previous work demostrated the induction of phosphorylation events as early as 2 min after substrate addition (Nguyen et al., 2016; Xiong et al., 2014). Many signaling pathways have already been implicated in the adjustment of fungal metabolism, which involve kinases or phosphatases.

The Mitogen-Activated Protein Kinase (MAP kinase) pathway, for example, represents a prototype of a signal distribution channel with phosphorylation being the major switch. In *N. crassa,* MAP kinases are known to be involved in cell-to-cell communication (Fischer et al., 2018), circadian regulation of cellular processes (Bennett et al., 2013; Lamb et al., 2012), phosphate signaling (Gras et al., 2013) as well as sensing of osmolarity and carbon (Huberman et al., 2017). Phosphorylation/dephosphorylation reactions involved in nutrient-sensing are furthermore central to the cyclic AMP (cAMP) signal transduction pathway (Aichinger et al., 1999; Ficarro et al., 2002; Lengeler et al., 2000). This pathway involves G-protein-coupled receptors (GPCRs), heterotrimeric G proteins, adenylate cyclase, cAMP-dependent protein kinases (PKA) (Lengeler et al., 2000; Lorenz et al., 2000; Xue et al., 1998) and phosphodiesterases. GPCRs are seven-helix transmembrane proteins anchored in the plasma membrane of eukaryotic cells, that are responsible for a diversity of functions, including environmental sensing, metabolism, immunity, growth, and development (Bock et al., 2014; Cabrera et al., 2015; Chini et al., 2013). However, how GPCR-mediated sugar sensing initiates the G-protein signaling reactions and then transduces this signal to the inside of the cell via heterotrimeric G proteins (Tesmer, 2010) is poorly understood. What is known is that cAMP, as the product of the adenylate cyclase, acts as an intracellular secondary messenger (Nauwelaers et al., 2006), regulating a variety of processes including plant cell wall degradation in *Trichoderma, Penicillium, Aspergillus* spp. and *N. crassa* (Dong et al., 1995; Farkas et al., 1990; Schuster et al., 2012) Low amounts of cAMP stimulate cellulase activity, whereas high levels of cAMP repress cellulase formation (Assis et al., 2015). It has been demonstrated in *N. crassa* that one Gα protein, GNA-1, associates with adenylate cyclase (CR-1) and is required for GTP-stimulatable activity (Kays et al., 2000). Some of the morphological aberrations of mutants in this pathway can be complemented by growth in the presence of exogenous cAMP (Kays et al., 2000; Kore-Eda et al., 1991).

Two-component regulatory systems, consisting of proteins with histidine kinase and/or response regulator domains are another system of widespread, ubiquitous signaling pathways in bacteria, slime molds, fungi and plants (Borkovich et al., 2004; Schaller et al., 2011). In fungi, all known histidine kinases are of the hybrid type (HHKs), containing both histidine kinase and response regulator domains within the same protein (Borkovich et al., 2004; Schaller et al., 2011). A histidine phosphotransfer protein (HPT) receives the phosphate from the response regulator domain of the HHK and in turn transfers this phosphate to an aspartate on a separate, terminal RR protein (Borkovich et al., 2004; Schaller et al., 2011). *N. crassa* possesses 11 HHKs, one HPT, two RRs (Borkovich et al., 2004) and one atypical RR, STK-12 (NCU07378) (Catlett et al., 2003; Park et al., 2011). Mutants lacking *stk-12* have defects in development of aerial hyphae during conidiation (Park et al., 2011). The 11 *N. crassa* HHKs contain additional domains at their N-termini required for normal hyphal growth, resistance to osmotic stress and fungicides, conidial integrity, female fertility (Alex et al., 1996; Schumacher et al., 1997), normal conidiation, pigmentation and perithecial development during the sexual cycle (Barba-ostria et al., 2011; Froehlich et al., 2005). Evidence suggests that the HHK OS-1 (NCU02815) and RRG-1 act upstream of the OS-2/HOG-1 MAP kinase pathway (Jones et al., 2007) and are required for proper control of OS-2 phosphorylation during the circadian rhythm (Vitalini et al., 2007). RRG-2 possesses a Heat Shock transcription Factor (HSF) DNA binding motif and is required for normal sensitivity to oxidative stress (Banno et al., 2007).

Differential phosphorylation of key components in substrate perception was demonstrated for the cellulolytic transcription factors CLR-1, a major cellulase regulator in *N. crassa* (Coradetti et al., 2012, 2013) and XLR-1 (a cellulase/hemicellulase regulator conserved in fungi (Samal et al., 2017) as well as a cellobionic acid transporter, CBT-1, after exposure to cellulose (Xiong et al., 2014). Carbon catabolite repression (CCR) is another central regulatory mechanism involved in substrate perception and metabolic regulation found in a wide range of microbial organisms that ensures the preferential utilization of glucose over less favorable carbon sources (Aro et al., 2005; Vinuselvi et al., 2012) and therefore affects the fungal response to nutrient availability (Fernandez et al., 2012; Nguyen et al., 2016; Bang Wang et al., 2017). Differential phosphorylation of CCR-related proteins after addition of different carbon sources was previously observed in a proteome analysis of *Trichoderma reesei* (Nguyen et al., 2016). In *T. reesei*, the phosphorylation of the transcription factor Cre1, a major mediator for CCR in fungi, at a serine in a conserved stretch within an acidic domain was furthermore found to regulate its DNA binding activity (Cziferszky et al., 2002).

Another protein class known to be involved in metabolic adjustments and CCR are F-Box proteins as part of the SCF complex (Skp, Cullin, F-box containing complex) (Assis et al., 2018; Colabardini et al., 2012; Jonkers and Rep, 2009; Skowyra et al., 1997). This multi-protein E3 ubiquitin ligase complex catalyzes the ubiquitination of proteins destined for the proteasomal degradation process (Assis et al., 2018; Welchman et al., 2005) and is, therefore, paving the way for new cellular or metabolic states. F-box proteins are found in all eukaryotes and display a large variety of functions. In fungi, they are involved in the control of the cell division cycle, meiosis (Krappmann et al., 2006), mitochondrial connectivity, control of the circadian clock components, fungal development (Kress et al., 2012), virulence (Jöhnk et al., 2016), glucose sensing, as well as the induction of cellulolytic genes (Borkovich et al., 2004; Colabardini et al., 2012). Phosphorylation reactions are a central element of F-box protein-mediated degradation, since F-box target proteins are commonly first phosphorylated before being recognized and ubiquitinated (Jonkers and Rep, 2009). For *A. nidulans,* Fbx23 and Fbx47 were identified as being important for CCR, with the former working in a ubiquitin ligase complex that appears responsible for CreA transcription factor degradation under carbon catabolite-derepressing conditions (Assis et al., 2018). In *T. reesei*, the ubiquitin C-terminal hydrolase CRE2 and the E3 ubiquitin ligase LIM1 play a role in the regulation of cellulase gene expression (Denton and Kelly, 2011; Glass et al., 2013; Gremel et al., 2008). However, the targets of the ubiquitin pathway have not yet been identified, nor is it clear whether the ubiquitinated factor is destined for degradation by the proteasome or if ubiquitination causes a signal-specific activation (Glass et al., 2013; Welchman et al., 2005).

Despite the importance and widespread occurrence of phospho-modifications as laid out above, the identification of sites of protein phosphorylation is still a challenge (Ficarro et al., 2002), hampering the elucidation of its role in global regulatory pathways, such as during substrate perception. Making use of phospho-proteomics analysis by state-of-the-art mass-spectrometry, we aimed to overcome this limitation and determine the phosphorylation patterns during the initial two minutes of contact with new carbon sources. We thus identified a large number of such post-translational modifications in proteins related to the recognition process of plant cell wall compounds by *N. crassa*, such as transcription factors, GPCRs, kinases and F-box proteins. Subsequently, *in silico* methods enabled us to construct putative proteome-wide protein-protein interaction (PPI) maps. Physiological analyses of several components within the cAMP signaling pathway that have clear substrate-dependent phosphorylation changes indicates that they are involved in metabolic adjustments to new carbon-conditions, including CCR.

Overall, this dataset will serve as a benchmark for identification and prediction of phosphorylation reactions during the sensing and signaling of cellulosic carbon sources in filamentous fungi.

## 2. Research material and methods

### 2.1 Cell culture and media shift experiments for proteome and phosphoproteome analysis

Flasks with *N. crassa* wild type strain (FGSC #2489) were pre-grown for 16 h on 2% sucrose medium, then washed three times with 1x Vogel’s salt solution (no carbon source added) for a total duration of thirty minutes before transferring to their respective carbon source: 1 mM galacturonic acid (D-GalA; Sigma Aldrich) plus 1 mM rhamnose (L-Rha; Sigma Aldrich), referred to as GalAR (D-GalA+L-Rha), 2 mM glucose (D-Glc; Sigma Aldrich), 2 mM xylose (D-Xyl; Sigma Aldrich), 0.5% cellobiose (Cel; Sigma Aldrich) and 1 mM 1,4-ß-D-glucosyl-D-mannose (referred to as Glucomannodextrins or Gm; Megazyme). Samples for global proteome and phosphoproteome were incubated for additional 2 min. No carbon condition (NC) was incubated for 1 h after the medium switch. More details about the experimental methodology for Mass Spectrometry can be found in the Supplementary Material.

### 2.2. Trypsin digestion

The samples were lysed, reduced, alkylated, and digested by trypsin. Digested samples were desalted using C18 solid phase extraction tubes (Supelco, St.Louis, MO, USA). The resulting peptide samples were concentrated and a bicinchoninic acid (BCA) assay (Thermo Scientific, Waltham, MA, USA) was performed to determine the peptide concentration and samples were diluted with nanopure water for MS analysis. 200 µg peptides aliquots for each sample were dried down for further IMAC enrichment, used for phosphoproteome analysis.

### 2.3 Phospho-peptide Enrichment by Immobilized Metal Affinity Chromatography (IMAC)

IMAC enrichment of phospho-peptides follows a previously established protocol (Mertins et al., 2018). Peptides were reconstituted in 80 % ACN/0.1% TFA and incubated with Fe^3+^-NTA agarose bead slurry. Then, the beads containing bound phospho-peptides were washed and desalted using the Stage Tips. After that, the phospho-peptides were eluted from the IMAC beads to the C18 membrane by washing the Stage Tip containing IMAC beads with 70 µl 500 mM phosphate buffer, pH 7.0 and 100 µl 1% FA before being eluted from the C18 membrane with 60 µl 50% ACN/0.1% FA. Eluted phospho-peptides were dried down and stored at -80°C until LC-MS/MS analysis.

### 2.4. Peptide and phopho-peptide identification by Mass-spectrometry based analysis

The peptide and phospho-peptide samples were both analyzed using a nanoLC system (Waters NanoAcquity LC, Waters Corporation) coupled to a Q Exactive™ Hybrid Quadrupole-Orbitrap™ Mass Spectrometer (Thermo Fisher Scientific) in data dependent mode. All mass spectrometry data were searched using MS-GF+ (Kim et al., 2008; Kim and Pevzner, 2014) to identify peptides by scoring MS/MS spectra against peptides derived from the whole protein sequence database. The MS-GF+ results were filtered based on 1 % false discovery rate (FDR) and less than 5-ppm mass accuracy. Based on spectral count MS data were considered only those peptides that were identified in at least 2 out of 4 experimental replicates with a positive spectrum, independent of the spectral count value. The phospho-peptide results were also searched using AScore algorithm (Gerber et al., 2006) to provide a probability-based score for each identified phospho-peptides. The mass spectrometry proteomics data have been deposited to the ProteomeXchange Consortium via the PRIDE partner repository (Perez-Riverol et al., 2019) with the dataset identifier PXD013964 and 10.6019/PXD013964.

### 2.5 Protein-Protein Interaction Networks

We constructed a system-wide network of the interactions between proteins found in the global proteome and phospho-proteome of *N. crassa* via prediction of functional interactions between the identified proteins, using the STRING database (Caccia et al., 2013; Szklarczyk et al., 2017). STRING database integrates and reassesses available experimental data from others curated databases, as well as information on structure, domains, function and similarity of proteins to predict functional protein–protein interactions or associations. In the STRING database, the predictions on protein-protein interactions are derived from the following sources: (i) systematic co-expression analysis, (ii) detection of shared selective signals across genomes, (iii) automated text-mining of the scientific literature and (iv) computational transfer of interaction knowledge between organisms based on gene orthology (Szklarczyk et al., 2017). While predicting interactions between *N. crassa* proteins using STRING database, we set the confidence score to be greater than 0.9 and the maximum of additional interactions to 10. Cytoscape was used for the visualization of the PPI network (Shannon et al., 2003). Supplementary Material 1 lists the proteins used to construct the PPI network.

### 2.6 Cell culture and F-box genes screening

The WT and deletion strains were obtained from the FGSC (http://www.fgsc.net) (McCluskey et al., 2010). The F-box genes were named following *A. nidulans* (Assis et al., 2018; Colabardini et al., 2012): *fbx-9* (*spp-1*, ΔNCU03658, FGSC#21255, Fbx9, AN10061), *fbx-19* (ΔNCU08642, FGSC#15478, Fbx19, AN4510), *fbx-20* (ΔNCU06250, FGSC#15629, Fbx20, AN4535) and *fbx-22* (*cdc4*, ΔNCU05939, FGSC#13190, Fbx22, AN5517). The corresponding deletion strain for *cr-1* (NCU08377) is FGSC#11514. *N. crassa* cultures were grown on Vogel’s minimal medium (VMM) (Vogel, 1956). Unless noted, 2% (w/v) sucrose was used as a carbon source. Strains were pre-grown on 3 ml VMM slants at 30°C dark for 24 hrs, then at 25°C in constant light for 4–10 days to stimulate conidia production. Conidia were inoculated into liquid media at 1 x 10^6^ conidia/ml in individual triplicates and grown at 30°C in constant light and shaking (200 rpm) in 24-deep-well plates. VMM media containing 2% D-Xyl (Sigma Aldrich) as control and 0.2% (w/v) 2-Deoxy-D-glucose (2-DG; Sigma Aldrich) plus 2% D-Xyl were used to test 2-DG influence, evaluated after 4 days. To test allyl alcohol sensitivity, 1% D-Glc +/-100 mM of allyl alcohol (Sigma Aldrich) were incubated for 2 days.

For xylanase activity assays, the strains were grown in 1% fructose VMM during 24 h at 30°C, then transferred to 2% D-Glc plus 1% D-Xyl VMM, or 1% D-Xyl VMM for 3 days. The xylanase activity was measured in the supernatant using Azo-xylan (from Beechwood; Megazyme) as substrate. Xylanase assays were conducted by mixing 25 µl of culture supernatant with 200 µl of Azo-xylan in 50 µl 1 M sodium acetate, pH 5.0, filling up to a final reaction volume of 400 µl with acetate buffer, and incubated at 37°C for 10 minutes. Reactions were stopped by centrifugation at 7000 x g for 2 min and by addition of 1 ml of precipitation solution (95% v/v ethanol) to the reaction supernatants. The absorbance was measured at 590 nm.

## 3. Results

### 3.1. A large number of proteins involved in signaling and perception are variant phosphorylated according brief exposure to new carbon sources

To identify the extent of initial protein phosphorylations upon exposure to new carbon sources, we performed switch experiments, in which sucrose pre-grown *N. crassa* was first starved for 30 minutes and then incubated for two minutes with mono- and disaccharides that are known to act as signaling molecules for plant cell wall-related polysaccharides. This short retention period was deliberately chosen to allow *N. crassa* to take up inducer molecules, but ensure that the proteome does not have time to significantly change, so that all measured differences are due to post-translational changes as a reaction to nutrient availability and not due to changes in enzyme abundances. A large number of proteins and phospho-peptides were detected by LC-MS/MS analysis in samples from these short-exposure experiments for each induction condition (Supplementary Material 2 – lists the peptides and phospho-peptides identified in different experimental conditions). We want to note that due to the presumed instability of histidine and aspartate phosphorylations during the enrichment strategy, we only searched for phosphorylation of serines (S), threonines (T) and tyrosines (Y) in our approach. We identified 2116 peptides that were phosphorylated equally in all conditions, but also between 207 and 506 phospho-peptides that were found to be phosphorylated specifically in only one condition (Figure 1A). We also studied the representation of the following functional classes among the proteins corresponding to these specific phospho-peptides: transcription factors (TFs), transporters, kinases, metabolism-associated proteins and proteins involved in signal transduction pathways (Figure 1B). The fact that a high number of these signaling-associated protein functions were present in the data sets (Table 1) strongly indicates that carbon sensing and signaling involves protein phosphorylation events as a central component in *N. crassa*.

**Figure 1:**
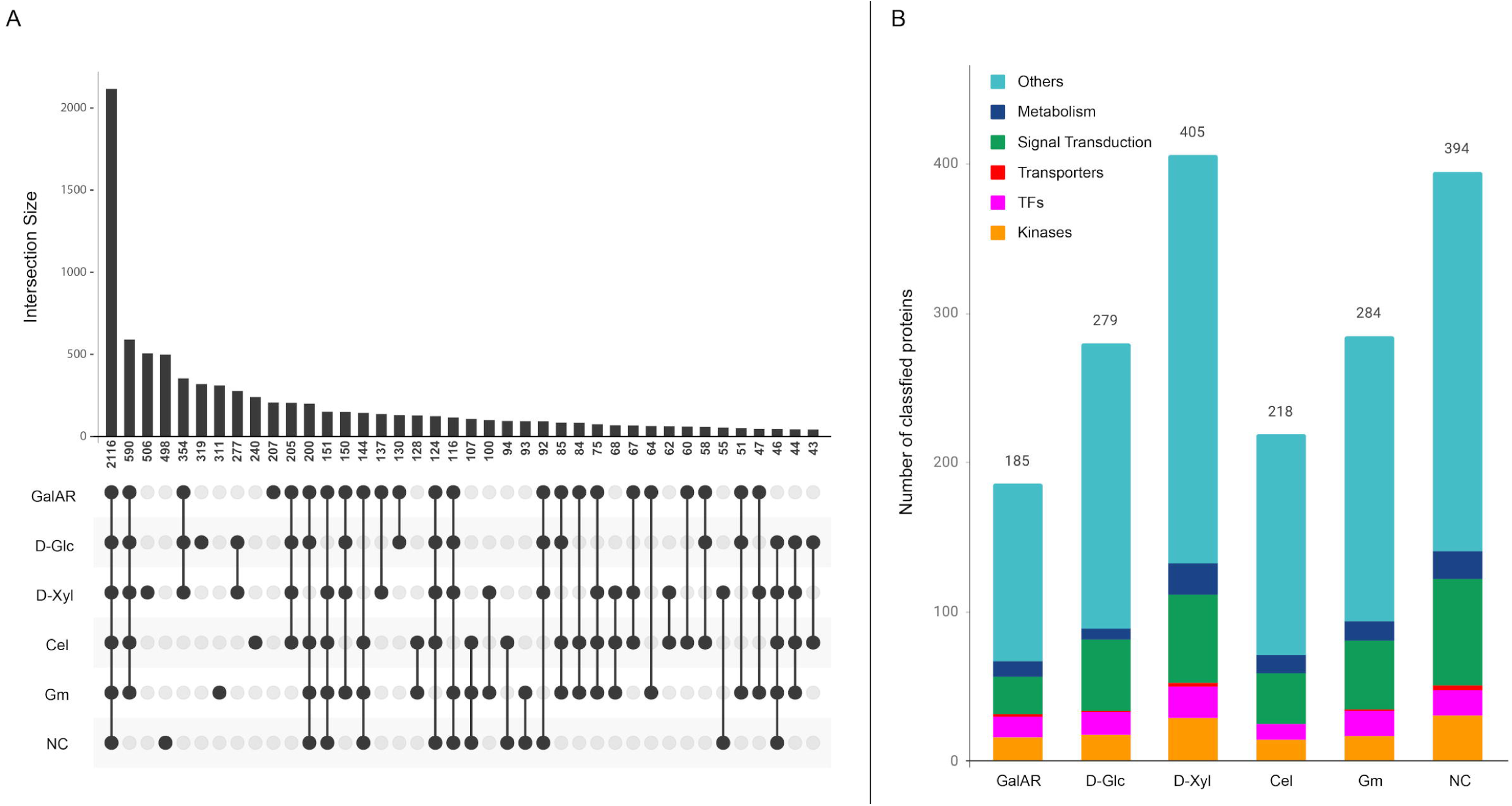
Proteome and phospho-proteome classification. A) Phospho-peptide distribution according to substrates. For this analysis, only the specific phospho-peptides (AScore >13, P<0.05) were considered. B) Functional classification of proteins with specific phospho-peptides for each condition. Substrate abbreviations: GalAR, D-GalA+L-Rha; D-Glc, D-Glucose; D-Xyl, D-Xylose; Cel, cellobiose; Gm, Glucomannodextrins; NC, no carbon.

**Table 1:**
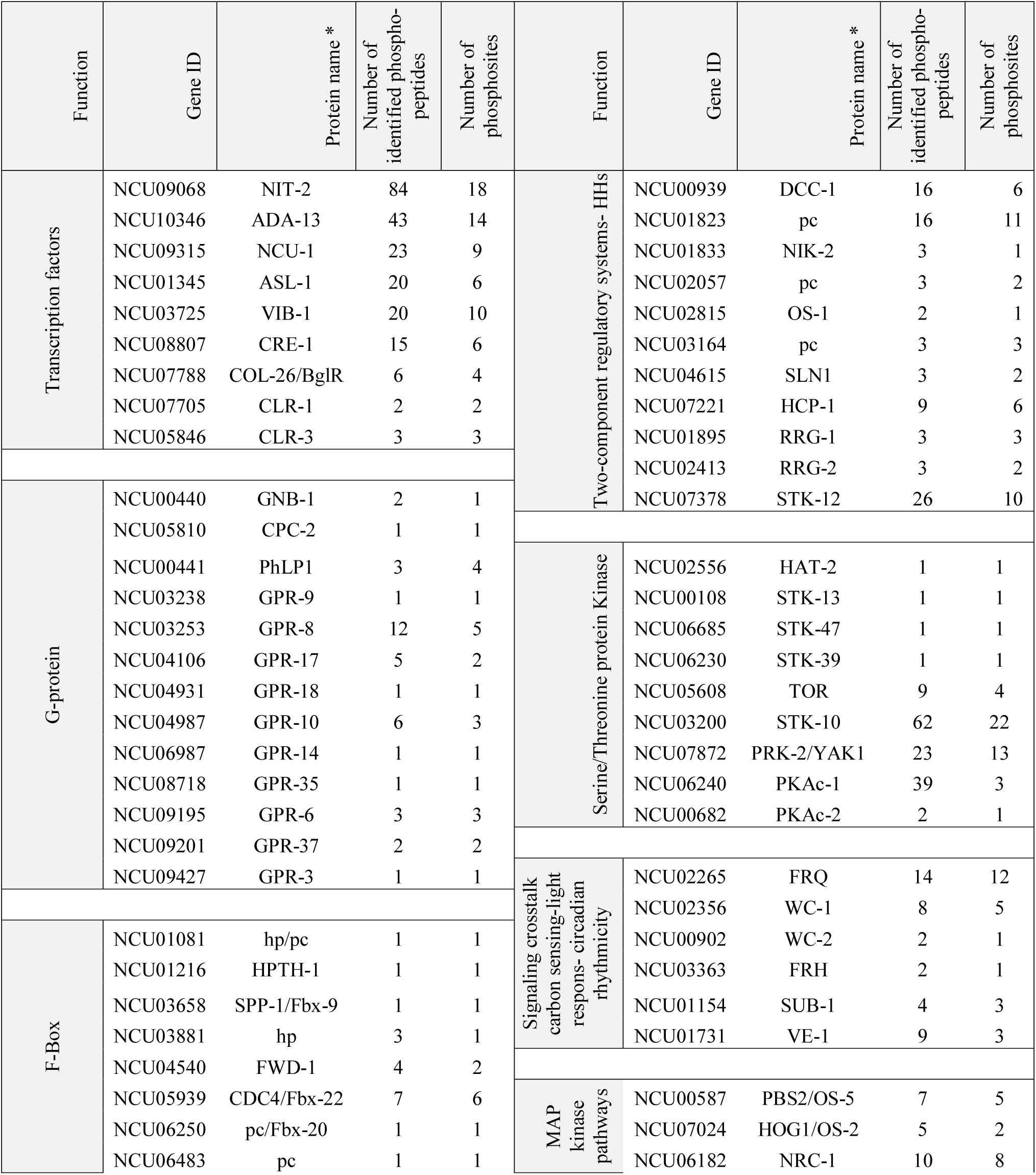
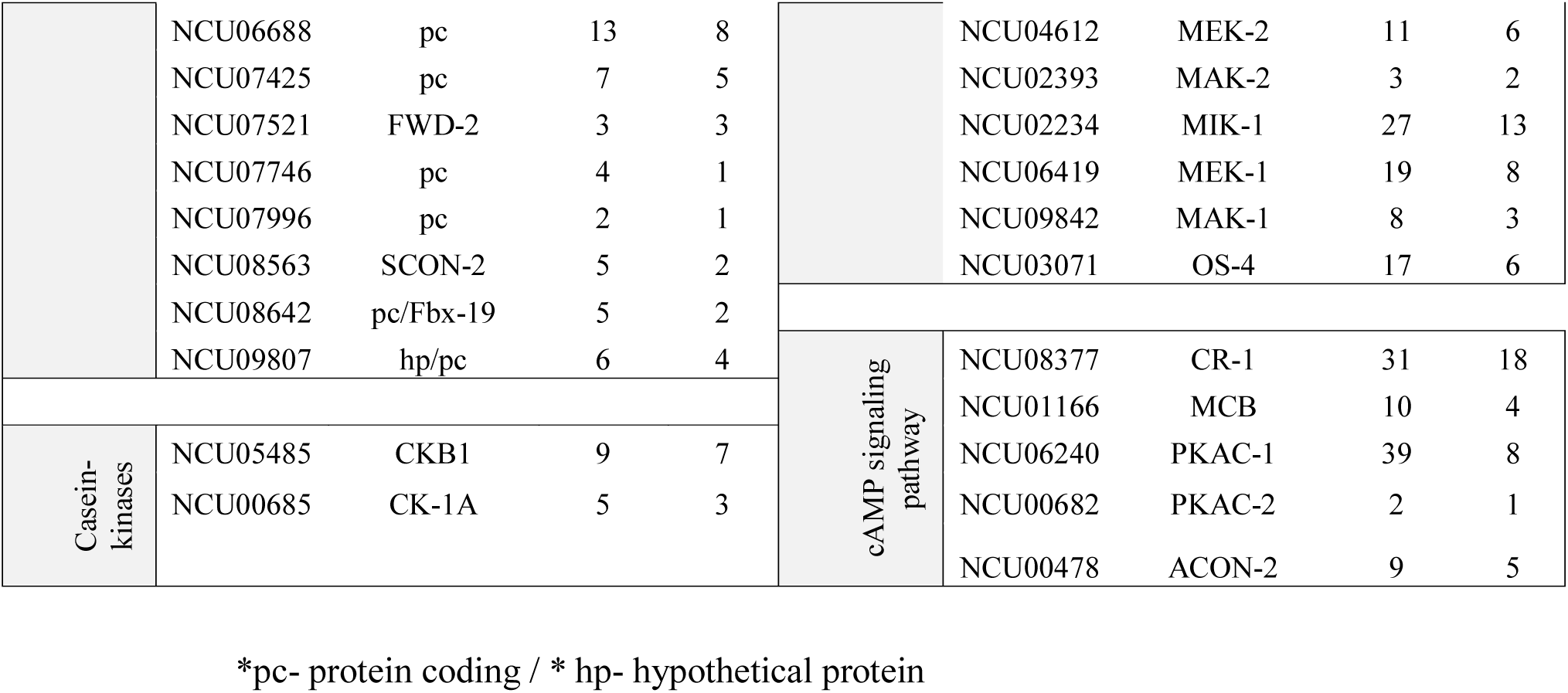
Summary of discussed classes of genes, with the respective protein name. The numbers of identified phospho-peptides correspond to all variant phospho-peptides identified experimentally. The number of phosphorylation sites correspond to the unambiguous identification of S, Y and T (Ascore > 13).

### 3.2 Protein-protein interaction networks predict changing signaling pathways in response to different carbon cues

A protein-protein interaction (PPI) network was constructed based on identified proteins in the proteomic and phospho-proteomic experiments, and Figure 2A visualizes this large-scale predicted network of interactions between *N. crassa* proteins present in the cell at the time point of the carbon switch. Thereafter we used the same functional classes (transcription factors, transporters, kinases, metabolism and signal transduction) to analyze the protein distribution across the network. The entire high-resolution network shows that proteins that belong to the same functional class have a tendency to cluster together (Supplementary Material 1). Nevertheless, complex interactions between proteins belonging to different functional classes are also observed in this network, especially in the central position of the network. To visualize the influence of substrates on this PPI network, the proteins that were identified been phosphorylated under each substrate were highlighted within the network (Figure 2B). Interestingly, it is possible to observe the differences in the profiles of phosphorylation for each substrate induction in Figure 2B. Specifically, the changes in distribution of highlighted protein nodes on specific induction conditions delineate the differential phosphorylation patterns that occurs in response to each carbon source substrate (Figure 2B).

**Figure 2:**
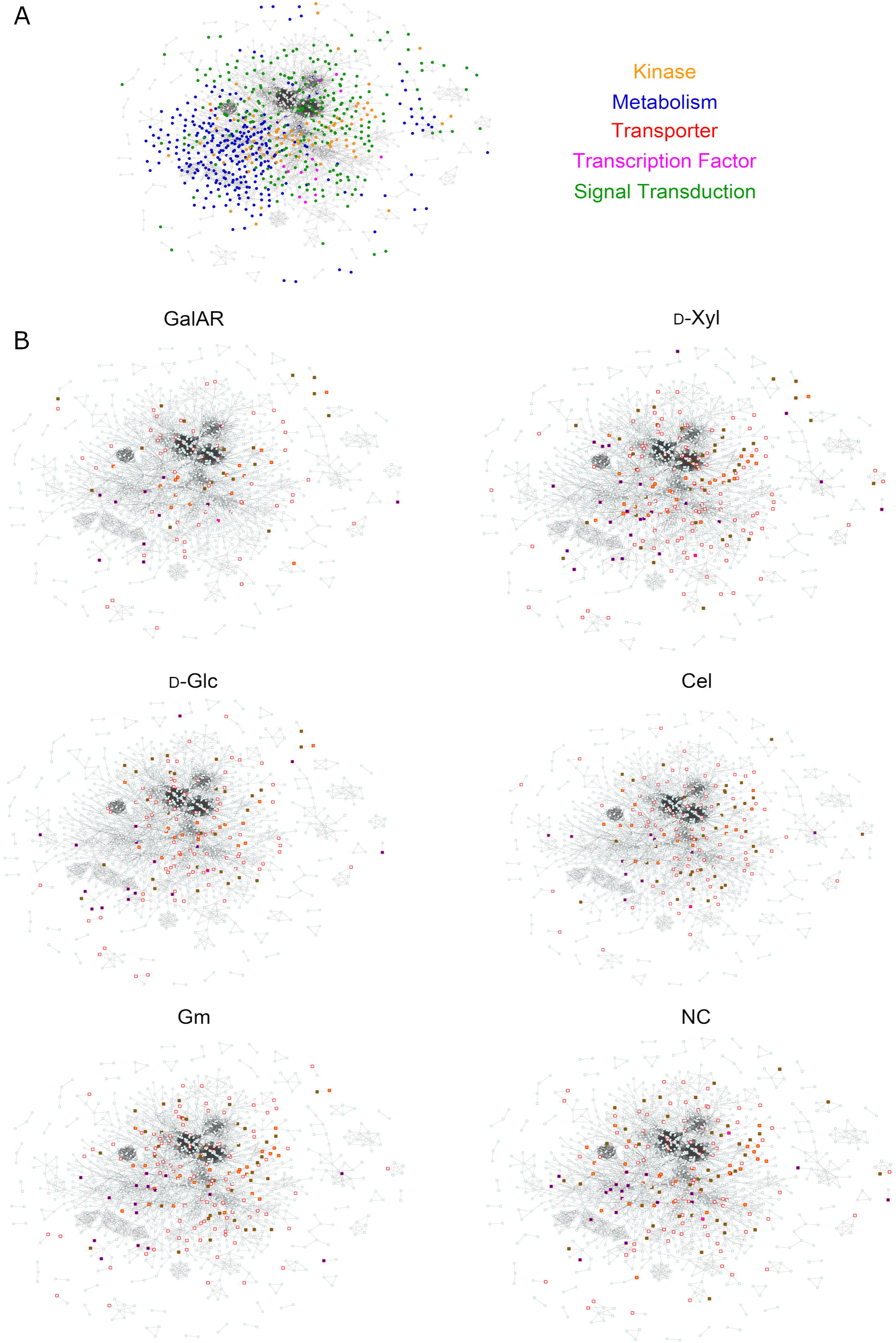
Protein-Protein Interaction network. A) The PPI network of all proteins identified by proteomics and phospho-proteomics in our experiments. Functional categories related to signaling are color-coded: green, signal transduction; orange, kinases; blue, metabolism; pink, transcription factors; red, transporters. B) Visualization of proteins in the PPI with specific phospho-peptides (independent of AScores) found in each experimental condition. Squares represent phosphorylated proteins, circles non-phosphorylated proteins. Red border-colored squares represent proteins phosphorylated specifically at the displayed condition and when filled, the proteins belong to one of the functional categories described in (A). GalAR (D-GalA+L-Rha), D-Glc (D-Glucose), D-Xyl (D-Xylose), Cel (cellobiose), Gm (Glucomannodextrins), NC (No carbon).

### 3.3 A large number of transcription factors exhibit differential phosphorylation upon carbon source perception

An earlier study had estimated the number of TFs in *Neurospora* to be 182 (Borkovich et al., 2004), a number that has been recognized as larger in recent analyses (Carrillo et al., 2017; Weirauch et al., 2014). Thus, we took a conservative list comprising 253 TFs included in the previous studies and inquired for the presence of phospho-sites in the corresponding proteins. One hundred-thirty TFs exhibited one or more phospho-peptides (Supplementary Material 2), with numbers going from one up to several dozens of modified residues per TF. Thus, for example, NIT-2 (NCU09068) exhibited a total of 84 different phospho-peptides, which represents a total of 18 *in vivo* unambiguously modified S and T (with Ascore >13) plus many more for which the localization is less certain. Of these, several are clustered in regions neighboring the GATA-type DNA binding domain (aa positions 742-792) including several that are phosphorylated in different residues. Determining which of the sites (many of which appear conserved in NIT2/AreaA homologues) could be playing regulatory roles, is an interesting question lying ahead.

Another TF exhibiting the second highest number of modifications is NCU10346 (ADA-13). This protein, with a predicted MYB DNA binding domain (PF00249), showed a total number of 43 phospho-peptides, comprising at least 14 unique phosphorylation sites.

Twenty-seven other TFs presented many identified phospho-peptides each, among which were NUC-1 (NCU09315), ASL-1 (NCU01345) and VIB-1 (NCU03725), all transcription factors important in mounting responses to changes in environmental conditions. Thus, for example VIB-1 has been associated with glucose sensing and carbon catabolite repression (Xiong, Sun, et al., 2014), playing a key role under cellulolytic conditions. The different VIB-1 phospho-peptides revealed 10 different phosphorylated residues, one of which also corresponds to a Y (Y107), the latter being modified only upon contact with D-GalA+L-Rha and Glucomannodextrins. T366, located less than 20 aa downstream from the VIB-1 DNA-binding domain, appears modified only in D-Xyl.

CRE-1 (NCU08807), the yeast Mig1 orthologue that plays a key role in carbon catabolite repression, showed six clearly identifiable phosphorylated residues (with Ascore >13) in four different regions of the protein (represented by five different peptides) including the extreme N-(S6, S11) and C-terminus. While four phospho-sites (S6, S11, S211 and T366) appeared modified in all conditions, some modifications were clearly condition-specific, such as S362 (only on cellobiose, Glucomannodextrins and without carbon source) and S209 (exclusively de-phosphorylated in cellobiose). Which phosphorylations may be modifying CRE-1 activity in *Neurospora* under different culture conditions remain to be determined.

COL-26/BglR (NCU07788) is a TF that has also been involved in glucose sensing/metabolism, regulation of starch degradation as well as the integration of carbon and nitrogen metabolism (Xiong et al., 2017; Xiong, Sun, et al., 2014). Six versions of two different phospho-peptides reveal four unique phosphorylation sites arranged in two pairs (S79/S83 and S674/S676). Interestingly, in glucose and D-GalA+L-Rha, all of the phospho-sites can be observed, while in no carbon condition only S676 was found to be modified.

The cellulase transcription factor CLR-1 (NCU07705) is phosphorylated at two sites, of which S59 phosphorylation was detected under all conditions tested. Phosphorylation of T120, which lies close to the Zn_2_-Cys_6_ binuclear cluster domain and within the predicted recognition sequence of a cAMP-dependent protein kinase (PKA) phosphorylation site, occurred specifically in the presence of Glucomannodextrins and D-Xyl. This latter phospho-site had already been detected in a previous study (next to S108, which was not identified here; (Xiong et al., 2014) to be more abundant on cellulose vs. no carbon, but found to be dispensable for activation. Of CLR-2 (NCU08042) and XLR-1 (NCU06971), which are also important regulators of cellulase gene transcription in *N. crassa*, no phospho-peptides were detected, peptides of this protein were also not found in the proteomics analysis, indicating a dominating transcriptional kind of regulation, as known for CLR-2 (Coradetti et al., 2012).

Although not a transcription factor, CLR-3 (NCU05846) is still highly relevant for cellulase regulation (Huberman et al., 2017) and was found to be phosphorylated in two regions and three potential sites (Supplementary Material 2). Interestingly, phosphorylation of T713 within a predicted casein kinase II phosphorylation site was specific for the absence of carbon.

### 3.4 Phosphorylations within two-component regulatory systems

Eight out of the 11 HHKs present in *N. crassa* (Borkovich et al., 2004) were found to be phosphorylated under at least one condition, including DCC-1/NCU00939, NCU01823, NIK-2/NCU01833, NCU02057, OS-1/NCU02815, NCU03164, NCU04615 and NCU07221 (Supplementary Material 2). Of note, the majority of these HHKs (5) were proteins with at least one PAS/PAC domain: DCC-1, NIK-2, NCU02057, NCU03164 and NCU07221. Four HHKs (DCC-1, NCU01823, NIK-2 and NCU07221) were phosphorylated under all conditions tested. NCU03164 was phosphorylated under every condition except D-Glc and D-Xyl. NCU04615 is unique in only being phosphorylated in the presence of Glucomannodextrins, while this is the only condition under which OS-1 and NCU02057 were not phosphorylated.

HPT-1 (NCU01489) was not phosphorylated under any conditions in this study. RRG-1 (NCU01895) was phosphorylated under all conditions, with three sites in total. RRG-2 (NCU02413) also had two phosphorylation sites and was not phosphorylated during growth on D-Xyl. The atypical RR STK-12 (NCU07378) had 12 regions of phosphorylation and was modified under all conditions. There were two different regions of phosphorylation on protein kinase domain (around S892, T893 and S903) and one in the AGC kinase, C-terminal domain (S1238) of STK-12.

### 3.5 Carbon specific phosphorylation in the MAP kinase pathways

The MAPKKK SskB/OS-4 (NCU03071), representing the first step in the osmolarity pathway, was found to be phosphorylated at S409 (and potentially S407) in response to D-Glc and Glucomannodextrins, with an additional ambiguous position for cellobiose. Phosphorylation of S1223, S153 and S77 was independent of the applied conditions. Response to cellobiose and no carbon resulted in phosphorylation of T1210, while D-Xyl as well as the D-GalA+L-Rha caused phosphorylation of S123. Interestingly, Glucomannodextrins and D-Xyl elicited phosphorylation of an SSTT-motif (S1343-T1346) within a potential PEST domain (Rechsteiner et al., 1989), suggesting a role for protein degradation in Glucomannodextrins and D-Xyl dependent carbon signaling. Although the AScore for these phosphorylation sites was below 13 (above the threshold of p=0.05; i.e. their precise localization within the peptide is not confirmed) phosphorylation of the respective peptides still indicates a relevance of the PEST domain modification.

In the MAPKK PBS2/OS-5 (NCU00587), condition-independent phosphorylation occurs at serines 219 and 346, while phoshorylation of S482 was only detected in the absence of carbon. The predicted PKC phosphorylation site at S63 is phosphorylated upon recognition of Glucomannodextrins, D-Glc and D-GalA+L-Rha. Phosphorylation of the serines at positions 77 and other sites adjacent to this was found to be condition-dependent for all conditions except no carbon indicating that this area may be a hot spot for post-translational regulation of carbon response in this kinase. HOG1/OS-2 (NCU07024), the MAP kinase of the osmolarity pathway, showed only two phosphorylated sites, one being a tyrosine. For Y173, phosphorylation was found under all conditions tested. However, phosphorylation of the adjacent T171 obviously required recognition of a carbon source, as no phosphorylated peptide was detected in the absence of carbon.

For NRC-1 (NCU06182), the MAPKKK of the pheromone response pathway, several phospho-peptides were detected. Six phosphorylated amino acids reside within predicted casein kinase phosphorylation sites and one in a PKC phosphorylation site. The serines at positions 49 and 558 were phosphorylated under all conditions tested. The predicted PKC phosphorylation site at S189 appears to be starvation-specific (no carbon) and T872 phosphorylation was only detected on cellobiose. In contrast to the osmolarity pathway, specific phosphorylations were also detected for D-GalA+L-Rha in combination with other carbon sources for NRC-1. In case of the MAPKK MEK-2 (NCU04612), we found phospho-peptides supporting the presence of both protein variants, with phosphorylation of T490 of variant 2 being present under all conditions tested. Despite detection of several phosphorylated sites, only few carbon-specific ones were found: while D-Xyl specifically elicited phosphorylation of S212, which is close to the active site, the serine at position 351 became phosphorylated upon sensing of D-Xyl, D-Glc and D-GalA+L-Rha. The MAP kinase of the pheromone response pathway, MAK-2 (NCU02393), only showed two phosphorylation sites, of which the tyrosine at position 182 was phosphorylated under all conditions tested. However, the adjacent T180 was phosphorylated only in the presence of D-Xyl and D-Glc.

In the cell integrity pathway, the MAPKKK MIK-1 (NCU02234) was found to be broadly phosphorylated under different conditions with up to 27 detected different phospho-peptides with 13 variant residues, albeit some have uncertain localization scores within peptides. In many cases, phosphorylation was not condition-dependent, but both for D-Glc (with S1215, a putative PKC site and S864, a putative casein kinase II site) and D-Xyl (with S1460, a putative cAMP-dependent site) specific sites were obvious. Phosphorylation of S450 was specific to the absence of carbon. Additionally, the region between S477 and S487 appeared to be strongly phosphorylated upon detection of carbon and may hence be a hotspot for carbon dependent regulation. Interestingly, this region overlaps with a poor PEST motif (Rechsteiner and Rogers, 1996), which hints at the involvement of protein degradation of MEK-1 (NCU06419) in regulation. Similarly, the region between amino acids 758 and 762 contains up to four phosphorylated sites specific for different conditions except “no carbon” and may as well represent such a hotspot.

In the MAPKK MEK-1, phospho-peptides were detected for different conditions, suggesting carbon-dependent phosphorylations also for this protein. Additionally, specific phosphorylations were detected for Glucomannodextrins on amino acids T92, S135, S139 and S184. The clustered serines at positions S130, S135 and S139 were phosphorylated in different combinations in various conditions and could thus represent an important area for carbon sensing. As also this region overlaps with a potential PEST motif, a contribution of protein stability or degradation in carbon sensing is further supported.

Phosphorylation of the MAP kinase MAK-1 (NCU09842) was found to occur on up to five sites, two of which being tyrosines. Thereby, phosphorylation of Y189 occured independently of the conditions tested, while Y185 was only phosphorylated in response to D-Glc (and potentially D-Xyl and Glucomannodextrins, but with low AScores). T187 became phosphorylated in response to plant cell wall-related carbon sources, but not in the absence of carbon.

### 3.6 Serine/threonine protein kinases are variably phosphorylated under different carbon conditions

Previous studies showed that there are 107 serine/threonine protein kinases in the *N. crassa* genome. According to our results, seventy-three out of the 107 serine/threonine protein kinases were found to be phosphorylated under at least one condition. It is worth noting that HAT-2 (NCU02556) is unique in only being phosphorylated in presence of cellobiose, implying that this modification may be important for cellulose utilization. Furthermore, STK-13 (NCU00108) and STK-47 (NCU06685) were only phosphorylated in response to Glucomannodextrins and D-GalA+L-Rha, respectively. Besides that, STK-39 (NCU06230) appears modified only in starvation condition.

In eukaryotes, protein kinase A (PKA) and the phosphatidylinositol 3-kinase TOR (NCU05608) are two major kinases involved in regulation of cell growth in response to nutritional signals. Yak1 and Sch9 are two downstream targets of these kinase cascades (Lv et al., 2015). Recent investigation showed that TrSch9 and TrYak1 play important roles in cellulase biosythesis in *T. reesei* (Lv et al., 2015). Our results demonstrated that STK-10 (NCU03200), the TrSch9 orthologue in *N. crassa*, exhibited the highest number of modifications upon recognition of cellobiose. This protein showed a total of 12 phosphorylated phospho-peptides (in 62 variants), which represents 22 *in vivo* unambiguously modified S, T and Y, suggesting that these sites may be playing regulatory roles. Similarly, PRK-2/YAK1 (NCU07872), the homolog of *T. reesei* TrYak1, showed 13 distinguishable phosphorylated residues (23 variants), represented in 12 different phospho-peptides under cellobiose condition. Thus, determining the function of these phosphorylation sites will be a worthwhile target for an increased scientific understanding of cellulase gene expression in filamentous fungi.

There are two catalytic subunits of PKA, namely PKAc-1 (NCU06240) and PKAc-2 (NCU00682), in *N. crassa*. Previous studies showed that PKAc-1 works as the major PKA in *N. crassa*, while deletion of *pkac-2* does not affect fungal morphology, suggesting that these two isoforms may play distinct roles (Banno et al., 2005). According to our results, PKAc-1 was found to be phosphorylated under all tested conditions, while PKAc-2 was phosphorylated only in the presence of D-Glc and cellobiose (for more details, see section 3.11.1).

### 3.7 Variances in phosphorylation of the G-protein-mediated signaling components

We analyzed the six G-protein subunits (Gα proteins GNA-1/NCU06493, GNA-2/NCU06729, GNA-3/NCU05206; Gβ proteins GNB-1/NCU00440 and CPC-2/NCU05810 and Gγ subunit GNG-1/NCU00041) (Muller et al., 1995; Yang et al., 2002), prosducin and 43 GPCRs, looking for phospho-sites in our dataset (Supplementary Material 2). Of the six G-proteins, only the Gβ subunits GNB-1 and CPC-2 (RACK1 homolog) were phosphorylated. GNB-1 was phosphorylated under all conditions in its extreme N-terminus, while S157 of CPC-2 was found to be phosphorylated only with Glucomannodextrins or without any carbon source.

The phosducin like-protein PhLP1 (NCU00441), which acts as a co-chaperone for β and γ subunits (Willardson and Tracy, 2012) is phosphorylated at S120 and S129 independent of the carbon source. The serines at position 41 and 47 are specifically phoshorylated in presence of Glucomannodextrins.

We furthermore detected 33 distinctive phospho-peptides for 10 GPCRs. This set contains NCU03238 (GPR-9), NCU03253 (GPR-8), NCU04106 (GPR-17), NCU04931 (GPR-18), NCU04987 (GPR-10), NCU06987 (GPR-14), NCU08718 (GPR-35), NCU09195 (GPR-6), NCU09201 (GPR-37) and NCU09427 (GPR-3). Of interest, four of these (GPR-17, 18, 35 and 37) are Pth11-related GPCRs, a large class previously implicated in sensing/degradation of cellulose in *N. crassa* (15). GPR-17 and GPR-35 were phosphorylated under all conditions, with an extra phospho-peptide observed only in the presence of D-Xyl for GPR-17 and during starvation (no carbon source) for GPR-35. GPR-9 was phosphorylated in D-Glc, D-GalA+L-Rha and D-Xyl. GPR-18 was phosphorylated in the presence of D-GalA+L-Rha, cellobiose and Glucomannodextrins, but not D-Glc or starvation, suggesting that this GPCR is phosphorylated only during growth on plant cell wall substrates. Results from RNAseq analysis showed that *gpr-18* is also among the most highly expressed of the Pth11-class GPCRs during growth on avicel (Cabrera et al., 2015).

The 10 GPCRs were variously phosphorylated in three different regions of the proteins. GPR-3, GPR-8, GPR-14, GPR-17, GPR-18, GPR-35 and GPR-37 are phosphorylated on residues in the intracellular C-terminal region. The mPR-Like/PAQR GPCRs GPR-9 and GPR-10 are phosphorylated on the first extracellular domain, while Stm1-like GPR-6 is phosphorylated in the large second intracellular loop region.

### 3.8 Signaling crosstalk between carbon sensing and components of light response and circadian rhythmicity

Even though *Neurospora* was grown under constant light conditions for the present study, the data reveal abundant phosphorylations present in clock components. Crosstalk between gene regulation in response to different nutritional conditions and the light response pathway is extensive (Schmoll, 2018; Stappler et al., 2017) and was also shown in *N. crassa* (Sancar et al., 2012; Schmoll et al., 2012). FREQUENCY (FRQ; NCU02265) is a key component of the *Neurospora* circadian clock, allowing precise daily control of several processes, including metabolism (Hurley et al., 2016; Montenegro-Montero et al., 2015). Under day-night regimes, FRQ is known to be subjected to extensive phosphorylations (> 75 unambiguous S/T sites) throughout its daily cycle, many of which are key in determining the proper pace of the clock (Baker et al., 2009; Larrondo et al., 2015; Tang et al., 2009). Herein, we have confirmed the phosphorylated status of 12 of these residues (seven with AScores >13), while identifying at least one additional site (T289). Out of the known sites, many appear to be phosphorylated under many conditions, while others appear to be modified only upon certain media transitions (Supplementary Material 2). Interestingly, the new *in vivo* determined phospho-site appears to be modified only under D-Glc. However, since we did not control tightly for sampling under identical time-of-day conditions, some of the observed changes might have been affected by the clock. Nevertheless, most of the sites observed in FRQ in this study have been shown to be *in vitro* substrates for CKI or/and CKII activity, such as S28, S145, T289, S462, S683, S685, and S950 (Tang et al., 2009).

Two other essential clock components, and also key actors in light-responses, are the transcription factors WC-1 (NCU02356) and WC-2 (NCU00902), which also impact regulation of plant cell wall degrading enzymes (Schmoll et al., 2012). WC-1 and WC-2 act as a heterodimer, forming the White Collar Complex (WCC). Our data verify discrete post-translational modifications of WC-2 in S433, a residue shown impact WCC activity and therefore circadian and light-responses (Sancar et al., 2009). Indeed, recently S433 was shown to be part of a P-cluster in WC-2 required for properly closing the circadian feedback loop (Wang et al., 2019). In the case of WC-1, three different regions were found to be phosphorylated, with one region (S1005-S1015) displaying multiple phosphorylations, particularly in D-Glc condition. These sites are located few residues distal from other phosphorylatable residues known to modulate circadian function of WC-1 (He et al., 2005; Wang et al., 2015; Wang et al., 2019) Some of the sites in WC-1 were recently shown to be phosporylable *in vitro* by CDK-1 (S831) or MAPK (S831 and S1015), without the need of a priming phosphorylation events (Wang et al., 2019). Interestingly, no phospho-peptides were detected for the LOV/PAS domain protein VVD (NCU03967) (Schafmeier and Diernfellner, 2011) which is involved in photoadaptation, despite the presence of several predicted phosphorylation sites and detection of the protein under all tested conditions. Finally, FRH (FRQ-interacting RNA helicase; NCU03363) is another key core-clock component, which *in vivo* studies have shown to display only one phosphorylation site (S21). Herein, we observed such phosphorylation in all analysed conditions.

The transcription factor SUB-1 (NCU01154) is predominantly known for its function in development in diverse fungi (Bazafkan et al., 2017; Colot et al., 2006), but has been shown to be connected to regulation of light response and chromatin remodelling by the white collar complex (Chen et al., 2009; Sancar et al., 2015). In SUB-1, we detected one unambiguous cellobiose-specific phosphorylation site (S381). S247 was phosphorylated under all conditions tested, while S143 was phosphorylated except upon recognition of cellobiose and in the absence of carbon, suggesting a starvation-triggered switch.

Also for VE-1 (Velvet; NCU01731), functions in development and secondary metabolism are best known (Bayram et al., 2008; Bayram and Braus, 2011), although this regulator also impacts plant cell wall degradation and light response (Aghcheh et al., 2014; Bayram and Braus, 2011). For VE-1 we detected one constant phospho-modification (S455) and two regions with potentially carbon sensing-relevant phosphorylations, one from T114 to S119, which overlaps with a poor PEST domain and a second one from T262 to T267. Interestingly, these regions contain two cellobiose-specific phosphorylations (T114 and T267, the latter with an Ascore only slightly below our threshold of 13.

The finding of clear condition-dependent phosphorylation patterns for both the photoreceptors and their regulators, such as SUB-1 or the casein kinases (see below) adds further support to the crosstalk between the reaction to carbon sources in the environment and sensing of light as well as circadian rhythmicity.

### 3.9 Phosphorylation of casein kinases

In *N. crassa*, casein kinases are mainly known for their function in the circadian clock (see above) with important impact on FRQ phosphorylation and stability (Diernfellner and Schafmeier, 2011), and are also know in other fungi (Al Quobaili and Montenarh, 2012; Apostolaki et al., 2012). *N. crassa* has one casein kinase II catalytic subunit and two casein kinase regulatory subunit as well as one casein kinase I (Borkovich et al., 2004).

The casein kinase II catalytic subunit alpha (CKA; NCU03124) did not display any detectable phosphorylation. In the casein kinase II regulatory subunit beta-1 (CKB1; NCU05485) however, S291 was found phosphorylated under all conditions tested, while S78, S101, S105, S106 and S223 show carbon-dependent phosphorylation specific to certain carbon sources (S101 and S105 for example to cellobiose and no carbon conditions). For the second casein kinase regulatory beta subunit (CKB2; NCU02754), we could not find any phosphorylated peptides, although the protein was detected under all tested conditions. The C-terminus of casein kinase I (CK-1A; NCU00685) on the other hand is phosphorylated at S313 and S320 (and potentially S312; low Ascore) independent of the condition tested, while S340 (and potentially S338; low Ascore) show carbon-specific phoshporylations including the absence of a carbon source.

### 3.10 Differential phosphorylation of F-Box proteins indicate that these modifications are necessary for correct protein turnover and metabolic adjustments

The genome of *N. crassa* is predicted to encode for 42 F-Box proteins. For 16 of these, we were able to detect phospho-peptides under at least one condition. Since homologs for some of these phosphorylated proteins had been implicated in regulatory functions of D-Glc utilization and CCR in *A. nidulans* (Assis et al., 2018), the respective *N. crassa* deletion strains were tested in more detail. First, we tested the capability of the strains to grow on different carbon sources (Figure 4A). Particularly the deletion strain for Fbx-22 (NCU05939; CDC4) displayed an aberrant growth phenotype, with improved growth, compared to WT, on D-Glc and glucomannan, while displaying reduced growth on arabinan, xylan and pectin. The deletion strain for Fbx-19 (NCU08642) displayed reduced growth on arabinan and pectin.

To test for a potential role in CCR, we next evaluated the resistance of WT and the mutants to 2-deoxy-glucose (2-DG) and allyl alcohol (AA) (Figure 4B). 2-DG is a non-metabolizable analogue of D-Glc that is often used to indicate impaired glucose repression in filamentous fungi (Allen et al., 1989; Assis et al., 2018; Xiong, Sun, et al., 2014). When functional, 2-DG is phosphorylated and activates CCR, resulting in the inability of the strain to grow on alternative carbon sources. When mutant strains are impaired in CCR, however, they become resistant to 2-DG. Of the four tested strains, the Δ*fbx-22* mutant showed some 2-DG resistance when 2% D-Xyl was used as an alternative carbon source in the presence of 2-DG (Figure 4B). AA is also used for assessing CCR, as reported for *M. oryzae* (Fernandez et al., 2012) and *A. nidulans* (Assis et al., 2018; Colabardini et al., 2012). When CCR is impaired, alcohol dehydrogenases are expressed despite the presence of D-Glc and will convert AA into the toxic compound acrolein. Thus, strains with impaired CCR exhibit AA sensitivity (Xiong, Sun, et al., 2014). As observed, the mutants Δ*fbx-9* and Δ*fbx-20* were insensitive, similar to WT. The mutants Δ*fbx-19* and Δ*fbx-22*, on the other hand, were sensitive to AA (Figure 4B). These data indicate that F-Box proteins Fbx-19 and particularly Fbx-22 might be involved in the correct signaling of, or adjustment to, conditions of high D-Glc concentrations.

The same AA-sensitivity with reduced growth and conidiation was also described for both homologous genes in *A. nidulans* (Assis et al., 2018). The deletion of these genes in the *A. nidulans* Δ*fbx19* and Δ*fbx22* strains furthermore led to reduced xylanase activity compared to the WT on D-Xyl (Figure 4C). In summary, these phenotypes strongly suggest an at least partially conserved defect of CCR control for Fbx-22 as well as potentially Fbx-19.

### 3.11 The adenylate cyclase CR-1 is extensively specific phosphorylated and acts as a central hub in the PPI

#### 3.11.1 CR-1 phosphosites identification

In the adenylate cyclase CR-1 (NCU08377), a central component of the cAMP signaling pathway known to be involved in nutritional sensing, we identified 18 unambiguos phosphorylated residues on 31 variant peptides (Figure 3A; Supplementary Material 2). Moreover, we found support for both isoforms of CR-1 with specific phosphorylation in both variants. CR-1 is heavily phosphorylated in both variants under different conditions. Phosphorylation of S67 was specific for cellobiose while T885 was specifically phosphorylated with D-GalA+L-Rha in variant 2. Glucose-specific phosphorylation only occurred in variant 1 at T191 and in the absence of carbon at S161 and S163 (only variant 1). For D-Xyl, specific phosphorylation occurred at T159 in variant 1 and around positions and in a different region at variant 2. Besides the support for two variants of adenylate cyclase by confirmation at the peptide level, these phosphorylation patterns indicate specific functions in carbon sensing and signaling of individual compounds for both variants. Consequently, *N. crassa* likely distinguishes these carbon sources and triggers specific responses also in terms of alternative splicing and coordinated phosphorylation for their utilization or degradation. Besides CR-1 as the core component, the cAMP pathway further comprises the functionalities of phosphodiesterases, which degrade cAMP in response to environmental signals, and protein kinase A (PKA; see also section 3.6). MCB (NCU01166), the regulatory subunit of PKA, showed phosphorylation in two regions of the N-terminus. In most cases, phosphorylation occured upon sensing of several C-sources or constitutively (e.g. T38, T40). However, phosphorylation of S41 was specific for recognition of D-Xyl and cellobiose. Phosphorylation of the catalytic subunit PKAC-1 (NCU06240) occured predominantly in two areas of the protein: from T19 to T32, containing eight putative phosphorylated sites, and from T377 to Y386. Both regions are overlapping with potential or poor PEST regions. In many cases, multiple parallel phosphorylations occured on several of the tested carbon sources. In contrast to PKAC-1, the second catalytic subunit of protein kinase A, PKAC-2 (NCU00682) showed only two phosphorylated amino acids. Albeit both having low AScores, phosphorylation of Y255 specifically occured in the presence of cellobiose and phosphorylation of T260 was specific for recognition of D-Glc.

**Figure 3:**
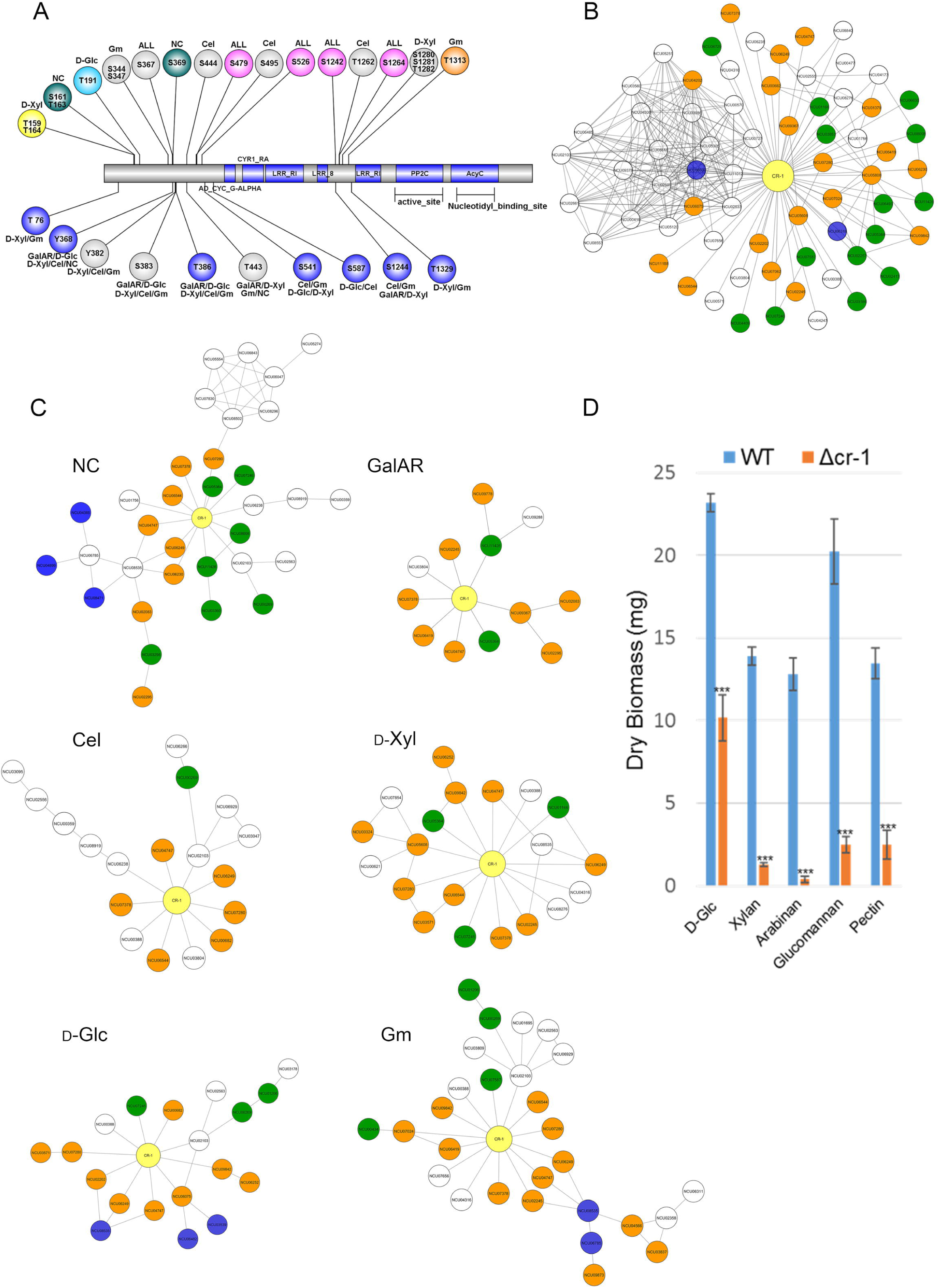
Carbon source-dependent changes of CR-1 interactions. A) Phospho-site distribution of CR-1 in response to substrate conditions. Picture shows all possible sites of phosphorylation. GalAR (D-GalA+L-Rha), D-Glc (D-Glucose), D-Xyl (D-Xylose), Cel (cellobiose), Gm (Glucomannodextrins), NC (No carbon). Colored circles represent sites with Ascore > 13, and grey circles with AScore < 13. B) Set of proteins directly interacting with CR-1. Colors according to functional categories: green, signal transduction; orange, kinases; blue, metabolism. C) Sub-PPI-networks of CR-1 nodes represent proteins that interact with CR-1, those peptides were found phosphorylated at specific condition. Colors according to functional categories described in (B). D) Mycelial biomass (dry weight) of the Δ*cr-1* mutant relative to on D-Glc, xylan, arabinan, glucomannan and pectin. Error bars represent the standard deviation from triplicate experiments. Statistical differences were calculated using *t*-test (***: p<10E-5).

**Figure 4:**
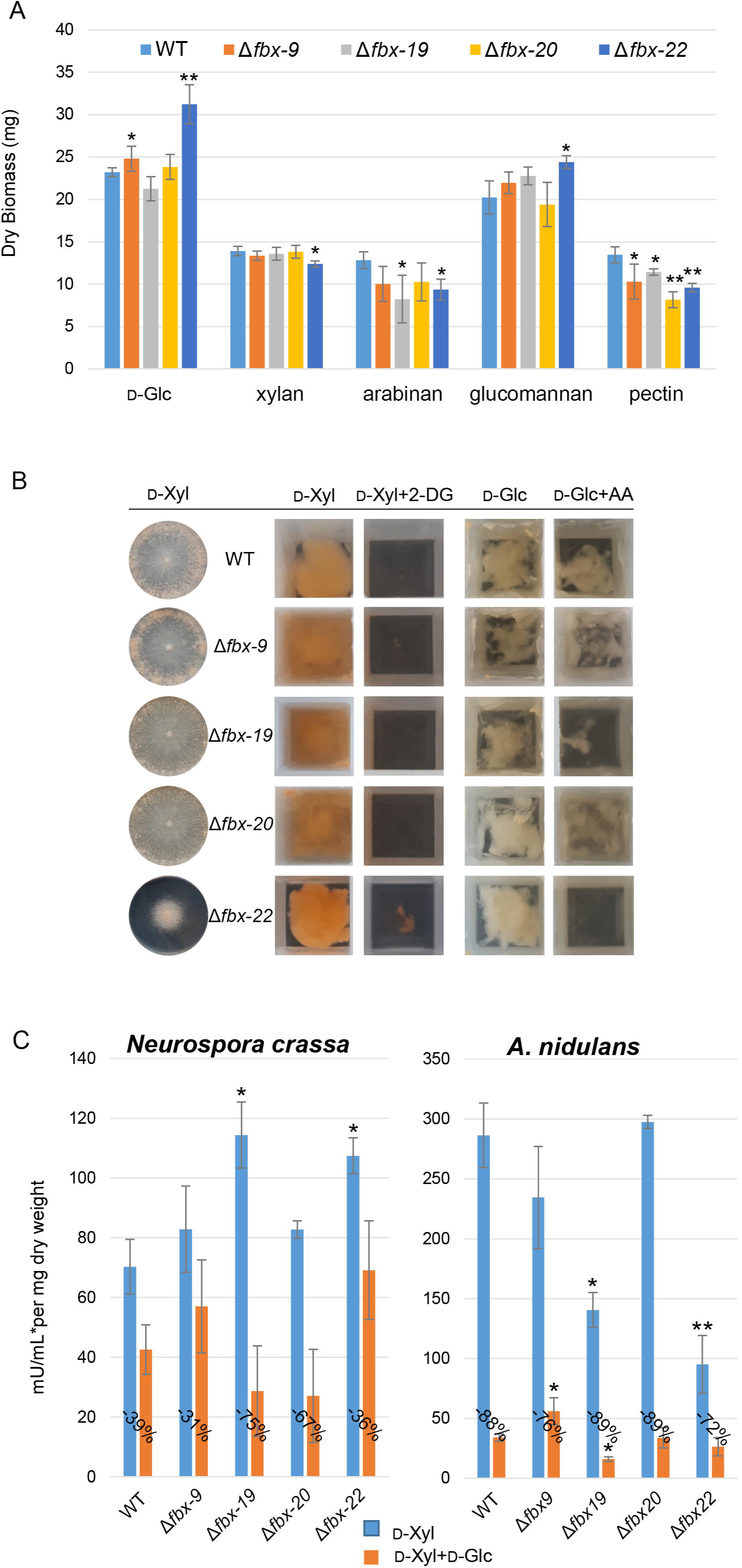
F-Box screening. A) Growth phenotypes of the F-box mutants on glucose (D-Glc), xylan, arabinan, glucomannan and pectin. B) The mutant strains were grown on 1% xylose (D-Xyl) plates. 2-DG resistance: deletion strains were grown on 2% D-Xyl as control and 2% D-Xyl + 0.2% 2-DG. Allyl alcohol (AA)–sensitivity: deletion strains were grown on 1% D-Glc as control and 1% D-Glc + 100 mM AA. C) Xylanase activities of F-box deletion strains of *N. crassa* and *A. nidulans*. Error bars represent standard deviation from triplicate experiments. Statistical differences were calculated between the mutants and WT within each treatment using *t-*test (*: p<0.05, **: p<0.001).

The high affinity phosphodiesterase ACON-2 (NCU00478) showed phosphorylation at several sites, in most cases relatively unspecific with respect to the carbon source (particularly S727 and S218). Nevertheless, carbon-specific phosphorylation was observed for Glucomannodextrins at T226 and D-Xyl as well as no carbon at S797, a site that overlaps with a predicted cAMP-dependent phosphorylation site. The low affinity phosphodiesterase PUR-1 (NCU00237) was not detected in the proteome.

#### 3.11.2 CR-1 interaction partners

Within the PPI, CR-1 is predicted to interact with 67 different proteins that form a complex PPI sub-network with a total of 265 connections (Figure 3B). For the visualization of potential carbon source-specific interactions around CR-1, we extracted from the entire network the phosphorylated CR-1-interacting proteins at each condition (Figure 3C). The sub-networks show the different possibilities of phosphorylation cascades that CR-1 might be involved in according to each induction condition. The predicted interactions are in line with the previously mentioned critical function of this gene in the regulation of growth and utilization of carbon sources (Kore-Eda et al., 1991; Lee et al., 1998). This is exemplified by enrichment of specific functional categories within the proteins of the CR-1 sub-networks (Ruepp et al., 2004) (Supplementary Material 3). For the sub-network from D-Xyl-induction, the most significantly enriched class was protein kinase (30.01.05.01; p-value 1.7E-16), while cAMP/cGMP-mediated signal transduction (30.01.09.07; p-value 7.7E-13) was dominating the Glucomannodextrins-induced subset (next to protein kinases, but with lower p-value: 1.0E-09). For the Cellobiose-induced subset, the top 7 classes were related to activities in the nucleus, such as DNA/RNA synthesis, modification and control (42.10.03, 42.10, 11.02.03, 10.01.09.05, 10.01.09, 11.02, 11.02.03.04, respectively; p-values 3.9E-13, 2.3E-12, 1.5E-11, 1.9E-11, 2.4E-11, 2.9E-11, 7.6E-11), followed by signaling pathways (30.01.09.07, 30.05.01.12.05, 30.05.01.12; p-values 7.7E-11, 3.6E-10, 1.5E-09). In the D-Glc sub-network, nucleus regulation, metal binding and serine/threonine kinase activity were overrepresented (42.10, 42.10.03, 16.17, 30.01.05.01.06; p-values 1.7E-10, 7.4E-10, 7.4E-10, 9.2E-10) plus the protein kinase class, cAMP/cGMP-mediated signal transduction and MAPKKK cascade (30.01.05.01, 30.01.09.07, 30.01.05.01.03; p-values 5.8E-9, 7.8E-9, 9.5E-9, indeed, a connection between MAPK and PKA pathways was described in *A. fumigatus* (Assis et al., 2015). The induction condition that seemed to trigger a high number of different pathways related to signaling and phosphorylation was D-GalA+L-Rha. In this condition, receptor-mediated signaling (30.05.01; p-value 5.5E-14), protein kinase (30.01.05.01; p-value 4.2E-13) and phosphate metabolism (01.04; p-value 3.43E-12) were dominating.

The analysis of the growth phenotype of the respective Δ*cr-1* deletion strain on different substrates demonstrated a substantially decreased capacity to degrade complex substrates (Figure 3D), further corroborating the role of CR-1 as a central hub in the cAMP signaling pathways involved in the sensing and response to environmental conditions.

## 4 Discussion

Numerous studies have indicated that rapid and reversible protein phosphorylations are involved in many perception pathways of environmental conditions across the tree of life (Cohen, 2000; Hornbeck et al., 2004; Lenoir et al., 2018; Nakagami et al., 2010; Sugiyama et al., 2008; Whitmarsh and Davis, 2000). Also in *N. crassa*, differential phosphorylation during growth on crystalline cellulose, sucrose and carbon-free medium has been described (Xiong et al., 2014). In the present study, we were specifically interested in the initial events of signaling at the onset of perception. For this purpose, we chose to study an extremely early time point (only two minutes after induction), which would be too short for substantial changes on either the transcriptome or proteome levels (Nguyen et al., 2016), but would give us an exclusive insight into rapid post-translational changes. To allow for a fast signal perception, we utilized soluble mono- and oligosaccharides that are known to act as inducing molecules in fungi, signaling the presence of the respective major plant cell wall polysaccharides: cellobiose for cellulose (Znameroski et al., 2012), D-Xyl for xylan (Sun et al., 2012), D-galacturonic acid and L-rhamnose for pectin (Thieme et al., 2017; Vries et al., 2002), and Glucomannodextrins for mannan (Ogawa et al., 2012). D-Glc and no carbon (starvation) were used as further controls. This extended set of induction conditions allowed us for the first time to identify signaling events that are specific for each of the cues derived from the individual components of a plant cell wall.

To better visualize the complex interplay of phosphorylation reactions, we constructed a protein-protein interaction (PPI) network with all detected proteins, exhibiting 12,066 predicted interactions based on available information gathered from STRING (Szklarczyk et al., 2017). The protein interaction prediction can infer physical and functional protein interactions from the genomic context (Browne et al., 2010; Shoemaker and Panchenko, 2007). PPI networks have already been used to provide insights into protein function and association with complex diseases (Kovács et al., 2019), fungal secretion pathways (Petranovic et al., 2013), identification and association of protein phenotypes (Hu et al., 2011) as well as domain interactions (Browne et al., 2010). Our PPI network represents the first view of possible *N. crassa* protein interactions at the early stage of substrate recognition. While the principal protein classes that putatively participate in the initial stages of substrate recognition (such as kinases, transcription factors, etc.) were found to be clustered across the network, numerous interactions between proteins of different functions are also represented. These interactions include many variances in phosphorylated proteins that can be hypothesized to be part of signaling pathways leading to metabolic adaptations. Detecting the vast changes of phosphorylation patterns across the network in response to different carbon sources strongly indicates that phosphorylation reactions are a central part of the environmental response of *N. crassa* that can be specifically adjusted to the composition of the substrate.

For transcription factors, kinases and other signaling components, such as CR-1, G-proteins and GPCRs, the phosphorylation profile is clearly affected by the substrate composition, providing novel insights into several well-known signaling pathways. For example, the differential phosphorylation of VIB-1, that acts upstream of CRE-1 (Xiong, Sun, et al., 2014), could be a mode of action how this TF is activated or inactivated and thus be involved in the modulation of the activity of CRE-1, potentially supporting the lignocellulolytic response. Further downstream in this regulatory cascade, the cellulase transcription factor CLR-1 shows condition-dependent phosphorylation as well. The regulatory impact of CLR-1 on the transcription factor CLR-2 (Coradetti et al., 2012) could be modulated by this phosphorylation and hence adjusted to the available substrate. As CLR-1 binds to its target promotors also under non-inducing conditions (Coradetti et al., 2012; Craig et al., 2015), the observed phosphorylation in presence of Glucomannodextrins and D-Xyl could help to keep CLR-1 de-activated in non-inducing situations.

To a large extent, our experiments enable the analysis of the phosphorylation reactions at amino acid resolution (reliably for all phospho-peptides with AScores >13, meaning a p-value for the phospho-site localization < 0.05). Regarding the G-proteins, RACK1-homologs have been demonstrated to function as an alternative Gβ in several fungi and are well-known for their additional role as scaffolding proteins for numerous processes in eukaryotes (Ron et al., 2013; Zhang et al., 2016). In *N. crassa*, the RACK1 homolog *cross-pathway control-2* (CPC-2) is required for proper expression of numerous amino acid biosynthetic enzymes in response to starvation for a single amino acid (Muller et al., 1995). Of interest, one of two residues that are phosphorylated by the WNK8 kinase in plant RACK1 proteins is T162 in the fourth WD40 repeat (J. Chen, 2015; Urano et al., 2015), just three residues away from the *N. crassa* phospho-site, which is also found during carbon starvation (no carbon condition). Genetic evidence suggests that phosphorylation of RACK1 on S122 and T162 negatively regulates its functions (Urano et al., 2015).

In terms of Gβ, studies in mammals have shown that morphine stimulation leads to phosphorylation and adenylate cyclase activity (Chakrabarti and Gintzler, 2003). Although the causative phospho-site has not been identified, phospho-proteomics studies have identified phosphorylated S residues in the N-terminus at positions 2, 72 and 74 (Chakravorty and Assmann, 2018), analogous to the N-terminal phosphorylation observed in this study. In *S. cerevisiae*, the Gβ Ste4p is phosphorylated, but on a T and S residue near the C-terminus of the protein (Chakravorty and Assmann, 2018). This region is not conserved in Gβ subunits from mammals or plants, and yeast mutants lacking both phosphorylation sites have relatively subtle defects in cell polarization during the pheromone response (Chakravorty and Assmann, 2018). In the crystal structure of the mammalian Gαβγ heterotrimer, the N-termini of the Gβ and Gγ form interacting α-helices that are important for dimerization (Moreira, 2014). Thus, it is intriguing that Gβ proteins from both *N. crassa* and mammals, but not *S. cerevisiae*, are phosphorylated in this region, perhaps influencing the association of the Gβ and Gγ subunits.

Also for the histidine kinase two-component systems, the identified phospho-sites within the domain structure of the proteins provide novel information that might be helpful to better understand their molecular setup. We found that a few of the *N. crassa* HHKs had at least one S/T phosphorylation site within a conserved motif in the protein. S/T phosphorylation of a plant HHK has been demonstrated for the cytokinin receptor AHK2 in *Arabidopsis thaliana* (Dautel et al., 2016; Nakagami et al., 2010; Sugiyama et al., 2008). One of these studies found that T4, S596 and T740 were phosphorylated, the latter two residues within the HK domain of AHK2 (Dautel et al., 2016). In *N. crassa*, two of the seven detected phospho-sites in DCC-1 (NCU00939) were identified within the RR domain, with S1338 and S1359 being phosphorylated under all conditions. NCU01823 was phosphorylated in different regions with two sites in the Orc1-like AAA ATPase domain, S722 and S723. Regarding NIK-2 (NCU01833), one phospho-peptide was detected at the extreme 3’ end of the first PAS/PAC domain being phosphorylated under all conditions. NCU03164 displayed four phosphorylation sites and is the only HHK with a site (S966) in the HK domain, being phosphorylated in presence of D-GalA+L-Rha, cellobiose, Glucomannodextrins and no carbon.

With the MAP kinase pathway, we evaluated a prototypical threefold phosphorylation cascade impacting osmosensing, pheromone response and cell wall integrity. According to Huberman et al., 2017, a relevant connection between carbon sensing and osmolarity pathway was described in *N. crassa*. The osmosensing (OS) MAPK pathway has been shown to act as a negative regulator of cellulase production in *N. crassa*, with the suggestion that osmolarity serves as a proxy for soluble sugars (Huberman et al., 2017). The Hog1 MAPK and several upstream components of the osmosensing signaling cascade are also required for normal expression of cellulases in *Trichoderma* (Wang et al., 2018). We found changes in phosphorylation across the whole MAP kinase pathway, with particularly heavy phosphorylation in the MAP KKK proteins. In contrast, MAP kinases HOG1, FUS3 and SLT2 contained only few phosphorylated sites. Carbon source-specific phosphorylation was detected for all the pathways, indicating that carbon sensing involves modification of all three MAP kinase pathways. Moreover, for several regions with increased phosphorylation frequency, we detected overlaps with PEST domains (Rechsteiner et al., 1996). Consequently, not only specific phosphorylation, but also protein degradation is likely to play a role in carbon sensing via the MAP kinase pathways. In many cases phospho-peptides with one phosphorylation or two phosphorylations were detected for MAP kinases, while multiple phosphorylation sites occurred in MAPKK and MAPKKK proteins. Hence, the conditions were selected correctly, i. e. during the process of signal recognition, in which the cascade is postulated to mediate the phosphorylation through the cascade from the MAPKKK (abundant phoshorylation) to the MAP kinase (moderate phosphorylation).

Another important class of proteins that also seems under regulation by phosphorylation are the F-Box proteins. Normally, F-box proteins are substrate receptors that recruit phosphorylated substrates, whereas phosphorylation of the F-box protein itself (due to lack of substrate) often results in autoubiquitination (by the SCF) and self-desctruction. However, phosphorylation of fungal F-box proteins can also affect other cellular functions, such as localization (Jöhnk et al., 2016). This study reveals specific phosphorylation of Fbx-22 (CDC-4) and a strong phenotype in the respective deletion strain, indicating that this gene has a critical function in CCR regulation that appears partly conserved in *N. crassa* and *A. nidulans.* The visualization of the Fbx-22 interaction network predicts that the protein might interact with two different clusters of proteins depending on the substrate conditions (Supplementary Figure 1). It remains to be determined how these interactions can influence the transcriptional response in the context of CCR regulation, showing a complex coordination of CCR by multiple signals.

The entire network visualization can show the differences in the phosphorylation behavior across the conditions, but cannot give a precise idea about the proteins involved in the interactions. We, therefore, decided to do a detailed analysis of the interactions of CR-1, since this enzyme is essential to the cAMP-dependent protein kinase pathway that is involved in cellulase and xylanase production (Assis et al., 2015). Especially for CR-1, the phosphorylation patterns changed drastically across the conditions. Interestingly, a major “hot spot” of phosphorylation appears to lie between two LRR (leucine-rich repeat) regions (Figure 3A). This motif is present in proteins with diverse functions and provides a versatile structural framework for the formation of protein-protein (Kobe and Deisenhofer, 1995; Kobe and Kajava, 2001) and ligand binding sites (Liu et al., 2017). As the structure of LRR motifs can lead to a conformational flexibility necessary to the protein interaction (Kobe and Deisenhofer, 1995), the phosphorylation in between these regions might also affect the binding properties by influencing the conformational structure of the binding sites. The phosphorylation of Ser/Thr residues between subdomains of LRR-RLK (leucine-rich repeat receptor-like protein kinase) was also previously described as being essential for catalytic activation of some protein kinases (Bojar et al., 2014; Liu et al., 2017).

Similarly intriguing, the identifiable sub-networks of CR-1 were found to have a specific design according to the interacting proteins with condition-specific phosphorylations. Extending the previously described importance of adenylate cyclase in development and growth (Ivey and Hoffman, 2005; Kore-Eda et al., 1991; Nauwelaers et al., 2006), our results therefore indicate that variances in phosphorylated CR-1 can interact with many other proteins and trigger specific phosphorylation cascades, depending on the composition of the medium.

## 5. Conclusions

The present study reveals fundamental changes in global protein phosphorylation during the initial moments of substrate recognition in *N. crassa*. The identification of protein phosphorylation followed by the prediction of protein-protein interactions allowed us to visualize the condition-specific phospho-protein distribution within the network. Many of the identified phospho-proteins are associated with sensing and signaling functions in the cell and thus appear to be involved in the regulatory networks governing the specific metabolic adaptation to plant cell wall polysaccharide degradation. Our protein-protein-interaction network analysis highlights that these differential patterns might lead to vast alterations in the respective sub-networks, as is the case for the adenylate cyclase CR-1, identified as a central hub with variances in phosphorylation patterns in the tested conditions. In many cases, the analysis provided amino acid level resolution, allowing to make assumptions about carbon source-specific structure-function relationships. Overall, the taken approach can therefore greatly help to identify novel functional interactions related to substrate signaling, and the experimental data can serve as a resource for future efforts to elucidate the underlying molecular mechanisms.

## Supporting information

Supplementary Methodology

## Authors’ contributions

MACH, NT, CT, LL, AS, and JPB jointly conceived the study, and each designed and supervised substantial parts of it. MACH, NT, YG, KEB, CDN, MAG, MSL, and LJA performed the experiments and acquired the data. All authors helped to analyze and interpret the data. MACH, NT, KAB, MS, LL, LFL, and JPB drafted the manuscript, which was critically revised by LJA, CT, GHB, AS and GHG. All authors read and approved the final manuscript.

## Acknowledgements

MACH thanks Prof. Anete Pereira de Souza (UNICAMP, Brazil) and the Coordenação de Aperfeiçoamento de Pessoal de Nível Superior, Computational Biology Program (CAPES Foundation, Brazil) for the post-doctoral fellowship (88881.161048/2017-01). GHG acknowledges support by the Fundação de Amparo a Pesquisa do Estado de São Paulo (FAPESP, Brazil), the Conselho Nacional de Desenvolvimento Científico e Tecnológico (CNPq, Brazil), and the Technical University of Munich – Institute for Advanced Study (TUM-IAS), funded by the German Excellence Initiative. LFL thanks for support by iBio, Iniciativa Científica Milenio (MINECON, Chile), CONICYT/FONDECYT (Chile) and the Howard Hughes Medical Institute (USA).

We are grateful for excellent technical assistance by Petra Arnold and Sabrina Paulus (TUM) and we would like to thank N. Louise Glass (UC Berkeley, LBNL) and Qian Liu (TIB, China) for critical reading of the manuscript and Yuxin Yhang (TUM, Germany) for the assistance with the graphical formatting. We furthermore want to acknowledge the Fungal Genetics Stock Center (FGSC; Manhattan, Kansas, USA) for providing their services and strain collection.

A portion of this research was performed using Environmental Molecular Sciences Laboratory (EMSL), a national scientific user facility sponsored by the DOE’s Office of Biological and Environmental Research and located at Pacific Northwest National Laboratory (PNNL). PNNL is a multi-program national laboratory operated by Battelle for the Department of Energy (DOE) under Contract DE-AC05-76RLO 1830.

## Competing interests

The authors declare that they have no competing interests.

## Funding

JPB: DFG grant # BE 6069/3-1

LFL: CONICYT/FONDECYT 1171151

LJA: 2014 / FAPESP, Brazil Grant numbers 2016/007870-9 and 2017/23624-0.

LL and CT: National Natural Science Foundation of China: 31761133018.

KAB acknowledges NIGMS GM068087 and GM086565 and NIFA Hatch Project #CA-R-PPA-6980-H.

