## Supplementary Methodology for "Broad substrate-specific phosphorylation events are associated with the initial stage of plant cell wall recognition in *Neurospora crassa*"

### Supplementary Methodology for Mass Spectrometry Analysis

*Reagents: Urea, dithiothreitol (DTT), iodoacetamide, iron chloride, Tris hydrochloride (Tris-HCl), trifluoroacetic acid (TFA), sodium deoxycholate (SDC), formic acid (FA), acetonitrile (ACN), methanol, phosphatase inhibitor cocktail 2 (PIC 2) and phosphatase inhibitor cocktail 3 (PIC 3) were obtained from Sigma (St. Louis, MO). The Ni-NTA agarose beads were obtained from Qiagen (Valencia, CA) and EmporeTM C18 extraction disks were from 3M (St. Paul, MN).*

2.2. Trypsin digestion

TissueLyser II system (Qiagen, Valencia, CA) trays were frozen at -20°C overnight. Two 3mm stainless steel beads were added to each sample tube and placed in the TissueLyser. The frozen samples were ground for 2 min at 30Hz until powderized and stored at -80 °C. A portion of the frozen powder was transferred into a 15 mL Falcon tubes containing 1 mL of lysis buffer (8M urea, 75 mM NaCl in 100mM NH4HCO3 pH 7.8, 10 mM NaF, 1% phosphatase inhibitor cocktail 2 (Sigma, P 5726) and 1% phosphatase inhibitor cocktail 3 (Sigma, P0044) on ice and vortexed into solution. A bicinchoninic acid (BCA) assay (Thermo Scientific, Waltham, MA USA) was performed to determine protein concentration at a 20x dilution. Following the assay, 5mM dithiothreitol (DTT) was added to the samples and incubated at 37°C for 1 hr with constant shaking at 800 rpm. Samples were then alkylated with 20 mM iodoacetemide for 1hr in the dark. The samples were then diluted 8-fold for preparation for digestion with 100 mM NH4HCO3, 1 mM CaCl2 and sequencing-grade modified porcine trypsin (Promega, Madison, WI) was added to all protein samples at a 1:50 (w/w) trypsin-to-protein ratio for 3 h at 37˚C. Digested samples were desalted using a 4-probe positive pressure Gilson GX-274 ASPEC™ system (Gilson Inc., Middleton, WI) with Discovery C18 100 mg/1 mL solid phase extraction tubes (Supelco, St.Louis, MO), using the following protocol: 3 mL of methanol was added for conditioning followed by 2 mL of 0.1% TFA in H2O. The samples were then loaded onto each column followed by 4mL of 95:5: H2O:ACN, 0.1% TFA. Samples were eluted with 1mL 80:20 ACN:H2O, 0.1% TFA. The samples were concentrated down to ~100µL using a Speed Vac and a final BCA was performed to determine the peptide concentration and samples were diluted with nanopure water for MS analysis. 200 µg peptides aliquots for each sample were dried down for further IMAC enrichment, used for phosphoproteome analysis.

#### 2.3 Phosphopeptide Enrichment by Immobilized Metal Affinity Chromatography (IMAC)

For IMAC, peptides were reconstituted at 1 µg/µl in 80% ACN/0.1%TFA prior to enrichment. The $Fe^{3+}$-NTA agarose beads were prepared by replacing the $Ni^{2+}$ ion on the Ni-NTA beads with $Fe^{3+}$ through the buffer exchange (1). Briefly, peptide samples were incubated with 10 µl 50% bead slurry at room temperature for 30 min with constant 1000 rpm shaking, spin down and the supernatant was discarded. The beads containing bound phosphopeptides were resuspended in 200 µl 80% ACN/0.1% TFA and loaded on Empore C18 silica-packed Stage Tips (2) for washing and desalting. Before sample loading, the Stage Tips were washed with 100 µl methanol, then twice with 50 µl 50%ACN/0.1%FA, and twice with 100 µl 1% FA. Beads were loaded on Stage Tip with pre-conditioned C18 membrane and were washed with 50 µl 80% ACN/0.1% TFA twice and 50 µl 1% FA once. After that, the phosphopeptides were eluted from the IMAC beads to the C18 membrane by washing the Stage Tip containing IMAC beads with 70 µl 500 mM phosphate buffer, pH 7.0 three times and twice with 100 µl 1%FA before being eluted from the C18 membrane with 60 µl 50%ACN/0.1% FA. Eluted phosphopeptides were dried down and stored at -80°C until LC-MS/MS analysis.

### 2.4. Peptide and phopho-peptide identification by Mass-spectrometry based analysis

The peptide and phosphopeptide samples were both analyzed using a nanoLC system (Waters NanoAcquity LC, Waters Corporation) coupled to a Q Exactive™ Hybrid Quadrupole-Orbitrap™ Mass Spectrometer (Thermo Fisher Scientifc). For phosphopeptide samples, the dried samples are reconstituted in 9 µl of 3% (vol/vol) MeCN/0.1% (vol/vol) FA right before LC-MS/MS analysis.

Both peptide and phosphopeptide samples were analayzed using an in-house packed 70-cm x 75 um i.d. 3-um Jupiter C18 column. Mobile phase A is 0.1% formic acid in water while mobile phase B is 0.1% formic acid in acetonitrile. The flow rate is 300nL/min. The gradient is as following: 0-2min 1% B; 2-20min 8% B; 20-75min 12%B; 75-97min 30%B; 97-100min 95%. For peptide analysis, a Q Exactive Plus in data dependent mode was used. Mass spectrometer settings were: full MS (AGC, 3 x $10^6$; resolution, 35000; m/z range, 300-1800; maximum ion time, 20 ms); MS/MS (AGC, 1 x $10^5$; resolution, 17500; m/z range, 200-2000; maximum ion time, 100 ms; minimum signal threshold, 5 x $10^4$; isolation width, 2 Da; dynamic exclusion time setting, 30 s; collision energy, 30). For phosphopeptide analysis, a Q Exactive HF in data dependent mode was used. Mass spectrometer settings were: full MS (AGC, 3 x $10^6$; resolution, 65000; m/z range, 400-2000; maximum ion time, 20 ms); MS/MS (AGC, 1 x $10^5$; resolution, 15000; m/z range, 200-2000; maximum ion time, 100 ms; minimum signal threshold, 5 x $10^4$; isolation width, 2 Da; dynamic exclusion time setting, 45 s; collision energy, 30).

All mass spectrometry data were searched using MS-GF+ software (3) (4) to identify peptides by scoring MS/MS spectra against peptides derived from the whole protein sequence database. The MS-GF+ results were then filtered based on 1% false discovery rate (FDR) and less than 5-ppm mass accuracy to generate a list of qualified peptide hit results. Only those peptides and phosphopeptides were considered for further analysis that were identified by positive spectral counts in at least 2 of the 4 biological replicates. The phsophopeptide results were also searched using AScore algorithm (5) to provide a probability-based score for each identified phosphopeptides.
