## Supplementary figures and images for "Broad substrate-specific phosphorylation events are associated with the initial stage of plant cell wall recognition in *Neurospora crassa*"

### Supplementary Figure 1

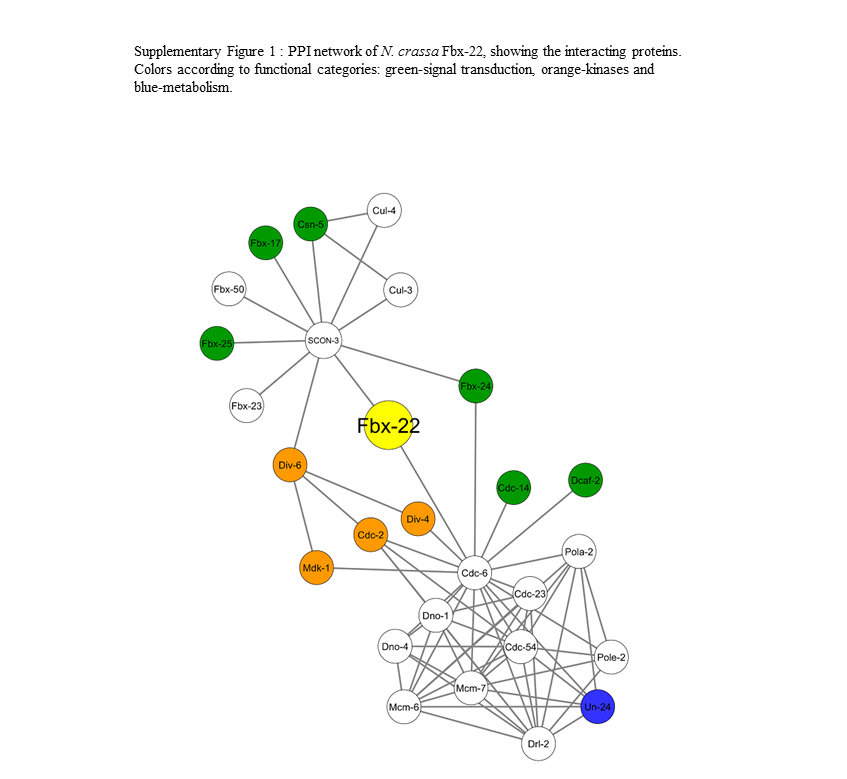
